# The conserved DNMT1 dependent methylation regions in human cells are vulnerable to environmental rotenone

**DOI:** 10.1101/798587

**Authors:** Dana M. Freeman, Dan Lou, Yanqiang Li, Suzanne N. Martos, Zhibin Wang

## Abstract

Allele-specific DNA methylation (ASM) describes genomic loci that maintain CpG methylation at only one inherited allele rather than having coordinated methylation across both alleles. The most prominent of these regions are germline ASMs (gASMs) that control the expression of imprinted genes in a parent of origin-dependent manner and are associated with disease. However, our recent report reveals numerous ASMs at non-imprinted genes. These non-germline ASMs are dependent on DNA methyltransferase 1 (DNMT1) and strikingly show the feature of random, switchable monoallelic methylation patterns in the mouse genome. The significance of these ASMs to human health has not been explored. Due to their shared allelicity with gASMs, herein, we propose that non-traditional ASMs are sensitive to exposures in association with human disease. We first explore their conservancy in the human genome. Our data show that our putative non-germline ASMs were in conserved regions of the human genome and located adjacent to genes vital for neuronal development and maturation. We next tested the hypothesized vulnerability of these regions by exposing human embryonic kidney cell HEK293 with the neurotoxicant rotenone for 24h. Indeed,14 genes adjacent to our identified regions were differentially expressed from RNA-sequencing. We analyzed the base-resolution methylation patterns of the predicted non-germline ASMs at two neurological genes, *HCN2* and *NEFM*, with potential to increase the risk of neurodegeneration. Both regions were significantly hypomethylated in response to rotenone. Our data indicate that non-germline ASMs seem conserved between mouse and human genomes, overlap important regulatory factor binding motifs, and regulate the expression of genes vital to neuronal function. These results support the notion that ASMs are sensitive to environmental factors and may alter the risk of neurological disease later in life by disrupting neuronal development.

## Introduction

DNA methylation refers to the addition of a methyl group (CH3) to the cytosine base of DNA by DNA methyltransferases. This predominately occurs at cytosine-guanine adjacent sites known as CpG sites. For most genomic loci, DNA methylation is coordinated across both inherited alleles. However, some loci maintain CpG methylation at only one allele and these regions are described to have allele-specific methylation (ASM; previously known as differentially methylated region DMR) (Bartolomei and Tilghman 1997). The most well-known of these regions are germline ASMs which control the expression of imprinted genes in a parent of origin-dependent manner. Imprinted genes are crucial in development and are commonly associated with genetic disorders such as Beckwith-Wiedemann, Angelman, and Prader-Willi syndromes (Butler 2009). In addition to the control of imprinted gene expression, DNA methylation is key to maintain genome stability via silencing retrotransposons (Chen et al., 2007; Walsh et al., 1998).

Investigations demonstrate that two types of genomic regions, imprinted germline ASMs and intracisternal A-particle (IAP)-like retrotransposons, seem vulnerable to environmental factors. Therefore, these two regions are proposed to be pivotal for understanding human disease in response to exposure and popularly pursued in animal and epidemiological studies (Jirtle and Skinner 2007; Murphy and Hoyo 2013). The former is attractive because exposure altered ASMs are anticipated to be faithfully transmitted to somatic cells during rounds of global demethylation and remethylation in early embryos (Barlow and Bartolomei 2014; Kacem and Feil 2009). As a result, parental exposure can be epigenetically inherited to modify offspring phenotype (Freeman and Wang 2019). The latter is exemplified in mice by the bisphenol A-hypomethylated IAP at the *agouti* gene for variations of coat color and obesity, as well as by altered methylation of IAPs at *Axin*^*Fu*^ for tail kinkiness (Rakyan et al. 2003; Zhou et al. 2007).

In our recent work, we developed two approaches, no-rescued DMRs (NORED) and methylation mosaicity analyses (MethylMosaic), to identify numerous genomic loci bearing potential ASMs (Martos et al. 2017). Both the known imprinted germline ASMs and newly identified ASMs are dependent on DNA methyltransferase 1 (DNMT1)(Li et al. 2015). Many of these novel ASMs are presumably sequence (single nucleotide polymorphism; SNP)-influenced ASMs (Kerkel et al. 2008). For example, in a reciprocal cross between 129S1/SvlmJ and Cast/EiJ or between C57BL/6NJ and Cast/EiJ, the Cast allele with SNP C of *Hcn2* ASM is always hypomethylated (i.e., independent of parental origin), whereas the 129 allele or the C57 allele with SNP A is always hypermethylated. Standing out of the previously appreciated sequence-dependent ASMs, a new paradigm of switchable ASMs that shows equal chances of either paternal or maternal allele to be methylated was revealed by our report (Martos et al. 2017). Importantly, the switchable feature seems also conserved in the human genome. At the *DLGAP2* locus, independent evidence confirms a maternally imprinted ASM during pre-implantation switched to a random ASM in somatic tissues during gestation (Monteagudo-Sánchez et al. 2018). Collectively, the mouse genome or human genome contains more ASMs (including both sequence-dependent and switchable ASMs) than previously appreciated (Deng et al. 2014, Martos et al. 2017, Monteagudo-Sanchez et al. 2018, Onuchic et al. 2018). The newly revealed random, switchable ASMs remind us of features in X chromosome inactivation, leading to a proposed hypothesis of regional autosomal chromosome inactivation (Wang et al. 2018).

Currently, germline ASMs are being increasingly considered in human disease; however, less studied are non-germline ASMs, which maintain CpG methylation at one allele independent of the parent of origin (Zhang et al. 2009, Deng et al. 2014, Martos et al. 2017). These regions regulate the expression of non-imprinted genes and these genes are hypothesized to have random monoallelic expression. Due to their predicted monoallelicity (DNA methylation and transcripts), we hypothesize that these regions are also targets for environmental factors and associated with disease like germline ASMs (Susiarjo et al. 2013).

The goal of this study was to determine whether our identified candidate ASMs in the mouse genome were in conserved regions of the human genome and to explore the possible adverse effects of differential methylation in these regions by examining their tissue expressivity and functional enrichment. Lastly, we tested our hypothesis that genes adjacent to non-germline ASMs would be vulnerable to environmental factors by exposing human embryonic kidney *HEK293* cells to the pesticide rotenone for 24h. We used whole transcriptome RNA-sequencing and targeted bisulfite-amplicon sequencing to evaluate changes in expression of genes and in DNA methylation at adjacent non-germline ASMs in response to rotenone. Indeed, our data demonstrate the vulnerability of these new, non-traditional ASMs to environmental exposure.

## Methods

### Identification of conserved DNMT1 dependent regions in the human genome

Whole Genome Bisulfite Sequencing (WGBS) was used to analyze base resolution methylomes of a series of *Dnmt1* (-/-), *Dnmt3a* (-/-), and *Dnmt3b* (-/-) murine embryonic stem cell lines (wild-type J1) as described previously (Li et al. 2015, Martos et al. 2017). DNMT1 dependent regions termed “NORED” were defined as regions with near complete loss of methylation in *Dnmt*1(-/-) compared to wild-type J1 that remained unable to recover methylation after the addition of exogenous *Dnmt1* cDNA. To identify the conserved DNMT1 dependent regions in the human genome, we used the UCSC Genome Browser LiftOver software to locate regions in the hg19 assembly from the mouse mm10 assembly. A text file of the chromosome positions (chr: start-end) for each putative ASM was uploaded into LiftOver and converted to the human hg19 assembly with a minimum ratio of 0.1 bases mapping for each region. The genomic location of each conserved region was analyzed in the UCSC Genome Browser window with NCBI RefSeq annotations. Transcription factor binding was observed using the Uniform Transcription Factor Binding data found in the ENCODE Regulation super track. We selected all transcription factor binding sites in H1-human embryonic stem cells (H1-hESCs) detected with CHIP-Seq experiments from the ENCODE consortium from 2007-2012 (ENCODE 2012).

### Functional enrichment for candidate DNMT1 dependent genes in the human genome

We restricted functional enrichment analysis of conserved human regions to those that had >70% base pair match for over 90% of the span of the identified region (Table 1). Pathway enrichment and network interactions for human genes nearest to these regions were calculated using the STRING database (Szklarczyk et al. 2014). Enriched cellular processes and molecular functions from Gene Ontology with FDR<0.05 calculated from the Fisher’s Exact test were reported. Network interactions were clustered using Cytoscape based on gene functional annotations in the reactome pathways (Fabregat et al. 2018).

**Table 1.**
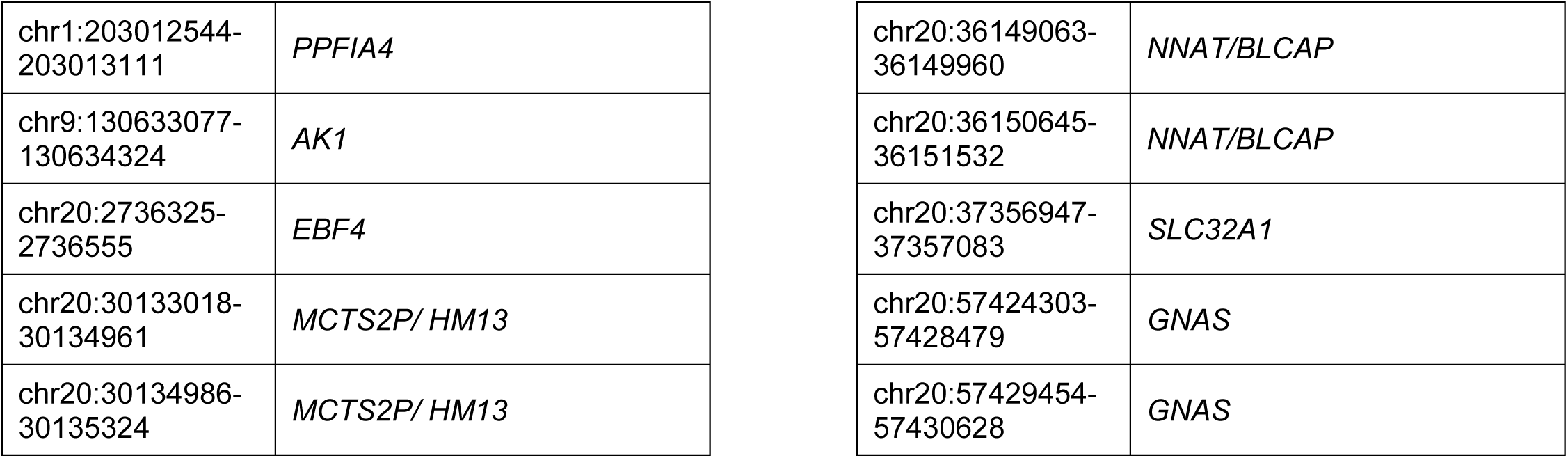
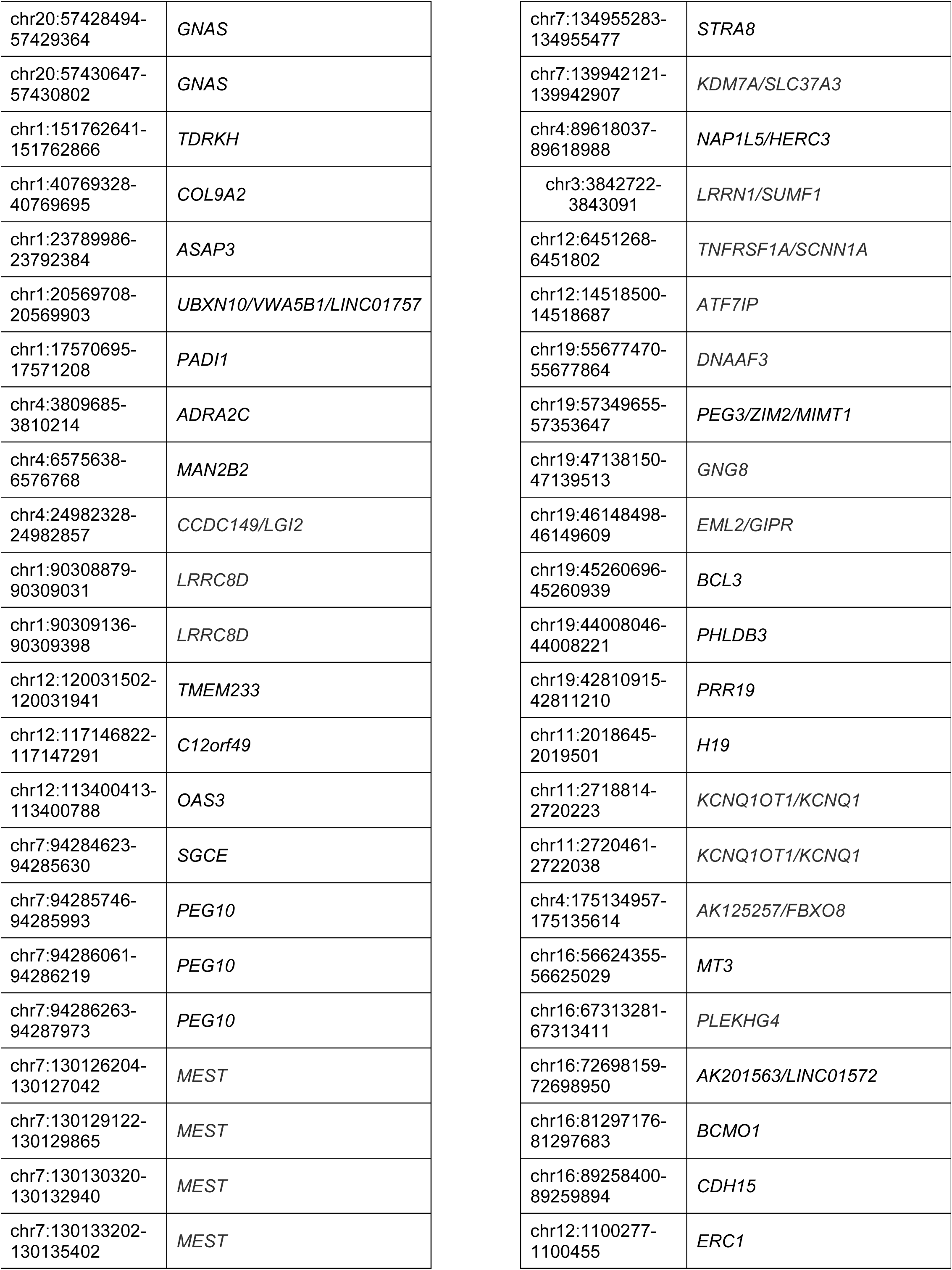
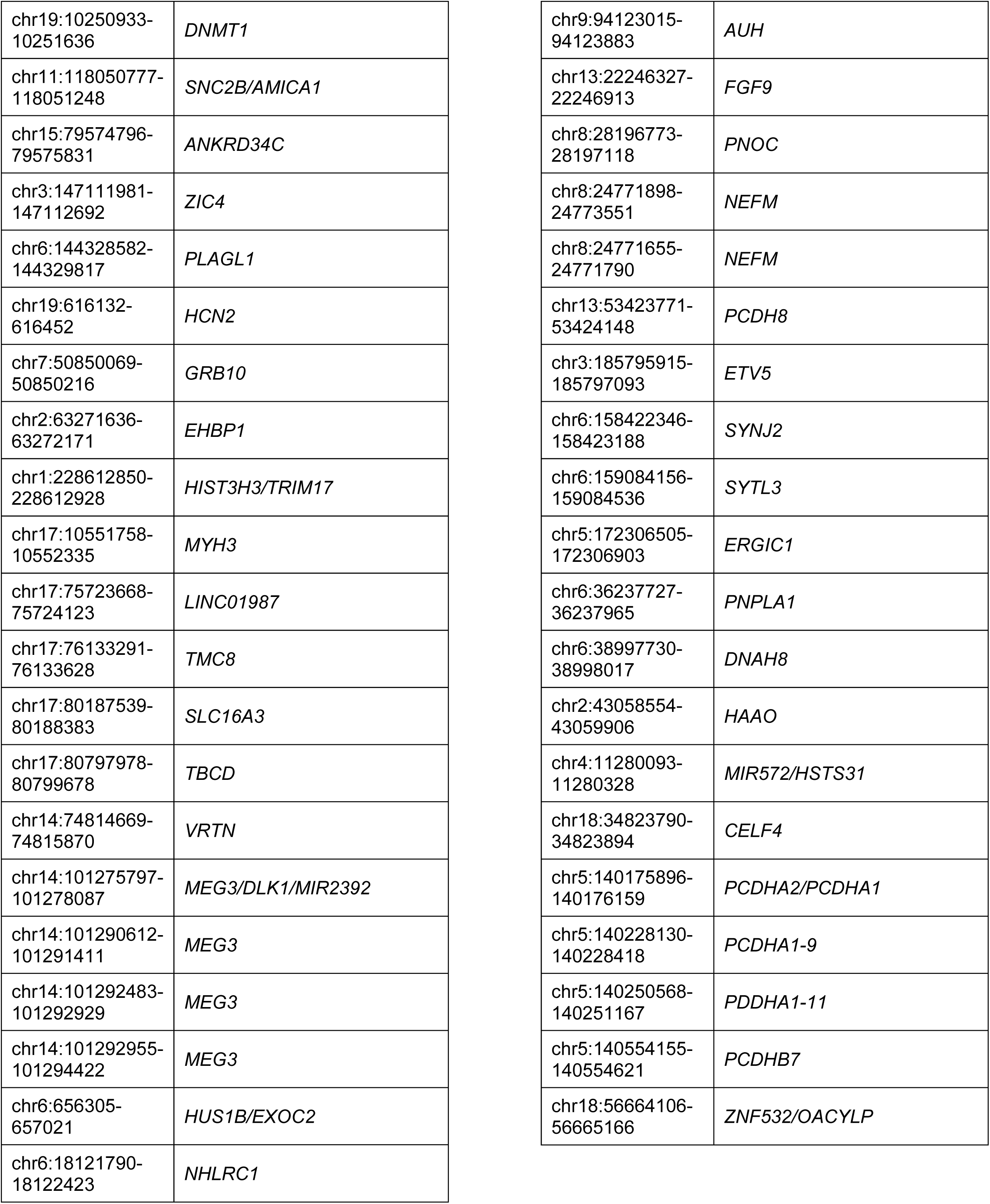
Predicted human DNMT1 dependent loci and genes.

### Tissue enrichment for candidate DNMT1 dependent genes in the human genome

The human DNMT1 dependent genes (Table 1) were used for tissue enrichment in EnrichR (Chen et al. 2013) using the ArchS4 database (Lachmann et al. 2018). The top six human tissues were reported with a p-value <0.01 adjusted using their correction for the fisher’s exact test (Chen et al. 2013). Enrichment of tissue-specific genes was performed using the TissueEnrich R package which uses gene expression data from the Human Protein Atlas (Jain and Tuteja 2018). Genes with an expression level of at least one transcript per million (TPM) were defined as tissue enriched when expression was at least five-fold higher in a distinct tissue compared to all tissues and tissue enhanced when expression was at least five-fold higher in a distinct tissue compared to the average expression of all other tissues. Group enriched genes were defined as genes with an expression level of at least one TPM and had at least five-fold higher expression in a group of tissues compared to all tissues. These definitions were taken from TissueEnrich.

### Cell culture and treatment of human cell line HEK293

All media reagents and chemicals in cell culture were purchased from Sigma (St. Louis, MO). Human cell line HEK293 was grown in Dulbecco’s Modified Eagle Medium with high glucose, L-glutamine, and sodium pyruvate. Media was supplemented with 10% (v/v) heat inactivated fetal bovine serum and 1% (v/v) Penicillin-Streptomycin. HEK293 cells were confirmed by ATCC. Cells were treated at approximately 70% confluency with 200nm rotenone or DMSO vehicle control (<0.001%) for 24 hours. Cell viability was measured with trypan blue (0.4%) staining. Viable and dead cells were counted manually using a hematocytometer. A minimum cell viability of 85% was used for all experiments.

### RNA extraction and RNA sequencing library construction

Total RNA was extracted from two replicates of DMSO or rotenone treated HEK293 using the trizol method (Invitrogen, Carlsbad, CA). A total of 2ug per sample was used for library construction using the TruSeq Sample Preparation kit from Ilumina (San Diego, CA). Poly-A containing mRNA molecules were isolated from total RNA using oligo-dT attached magnetic beads. Isolated mRNA was then fragmented and synthesized into double stranded cDNA according to kit instructions. Ligation of unique Ilumina adapter indices was completed for each sample before bead purification. Libraries were loaded onto a 2% agarose gel and library products between 200-800 bp were purified using the mini-Elute gel extraction kit from Qiagen (Hilden, Germany). Approximately 150 ng was sent for sequencing on a HiSeq 2000 platform with 100 bp paired-end reads.

### RNA sequencing data analysis

Adapter sequences were removed from the raw sequencing data and individual libraries were converted to the fastq format. Sequencing reads were aligned to the human genome (hg19) with TopHat2 (v2.0.9) (Kim et al. 2013). For mRNA analyses, the RefSeq database (Build 37.3) was chosen as the annotation references. Read counts of annotated genes were obtained by the Python software HTSeq-count (Anders et al. 2015). The read counts of each transcript were normalized to the length of the individual transcript and to the total mapped fragment counts in each sample and expressed as fragments per kilobase of exon per million fragments mapped (FPKM) of mRNAs in each sample. Differentially expressed genes were defined as those with a 1.5 fold change in expression using a FDR<0.05 from the edgeR package (Robinson et al. 2010). We examined differentially expressed genes in RNA-seq in common with the genes nearest to the conserved DNMT1 dependent regions in the human genome with >70% base pair match for over 90% of the span of the identified region in the mouse genome. Overlapping genes were entered into the Genotype-Tissue Expression (GTEx) database using the multi-gene query tool (GTEx Consortium 2013). Parkinson’s disease brain regions associated with motor function including the cerebellum, cortex, frontal cortex, spinal cord, substantia nigra, and basal ganglia were selected for further expression analysis.

### RNA sequencing validation with quantitative reverse transcription-PCR

Total RNA was extracted from an additional replicate of HEK293 treated with DMSO or rotenone using the same procedure as stated above. A total of 500ng RNA was converted to cDNA with the PrimeScript RT reagent kit with gDNA eraser from Takara (Kusatsu, Japan). We selected seven out of fourteen overlapping genes for quantitative PCR (qRT-PCR) analysis using primers listed in Supplemental Table 1. All qRT-PCR reactions were performed on a 7500 Real-Time PCR system from Applied Biosystems (Foster City, CA) using the iTaq Universal SYBR Green Supermix from Bio-Rad (Hercules, CA). The change in expression was normalized to the GAPDH housekeeping gene and expressed as fold change (2-^ΔΔCT^).

### Bisulfite-DNA conversion and Bisulfite-amplicon sequencing library construction

Genomic DNA was extracted from two replicates of DMSO or rotenone treated HEK293 using Phenol: Chloroform: Isoamyl alcohol (Sigma, St. Louis, MO). A total of 200 ng DNA was Bisulfite-converted using the Sigma DNA Imprint Modification kit two-step protocol. Bisulfite-converted DNA (BS-DNA) was amplified with primers for selected DNMT1 dependent regions designed with MethPrimer (Li and Dahiya 2002) (Supplemental Table 2). Amplified BS-DNA products were run on a 2% EtBr agarose gel and purified using the mini-Elute gel extraction kit from Qiagen (Hilden, Germany). Purified products for each sample were pooled together and 1 ng was used for library preparation using the Ilumina Nextera DNA Library Preparation kit. Each sample was tagged with a unique Nextera XT adapter (San Diego, CA). Sequencing libraries were quality checked via Bioanalyzer and run on an Ilumina MiSeq platform to generate 150 bp paired end reads.

### Bisulfite-amplicon sequencing analysis

The raw fastq files were imported into the Galaxy web platform (Afgan 2016). Reads with quality score >30 were trimmed with Trim Galore (Krueger 2015). Reads were mapped to amplified sequences in the human genome (hg19) using bwa-meth (Pederson et al. 2014). MethylDackel was used for methylation calling and per-cytosine contexts were merged into per-CPG metrics (https://github.com/dpryan79/MethylDackel). Duplicates and singletons identified in alignment were ignored from the methylation call. Minimum and maximum per-base depths were 1000x and 100,000x, respectively. The output was selected for methylKit format. Coverage statistics and differentially methylated regions were calculated for CpG sites with methylKit installed in R (v3.5) (Akalin et al. 2012). Differentially methylated cytosines were defined as being present in both biological replicates, having a minimum absolute difference of 1% using the coverage weighted mean, and having a SLIM adjusted q-value<0.01 using the methylKit logistic regression model (Ning et al. 2011). The change in mean percent methylation (Δme) for all CpG sites within a defined region was calculated by taking the mean number of methylated versus non-methylated CpG sites from the pooled control and treated samples and using Fisher’s exact test FDR <0.05.

## Results

### DNMT1 dependent regions in the mouse and human genome

Two approaches, NORED and MethylMosaic, independently identified over 2,000 regions. To simplify future interpretation, we focused on 207 overlapped regions (i.e., ‘NORED+MethylMosaic’ regions) to initiate our investigation. We compared these 207 regions from mouse and observed 145 of these regions were conserved in the human genome. Most regions identified in the human genome were highly conserved with >70% matched bases for more than 90% of the entire span of the region (Figure 1A-B). Analyzing these regions in the genome browser, we noted that 70% of the conserved regions in the human genome were located in the gene body and approximately 50% of them had transcription factor binding sites in human embryonic stem cells (Figure 1C-D). The two transcription factors that were the most significantly enriched were POL2RA and TAF1 with binding sites at 19% of conserved DNMT1 regions. Both were concentrated around transcription start sites and regulate RNA polymerase II binding and processivity in gene transcription. The third most enriched transcription factor was CTCF with binding motifs in 17% of conserved regions and most often found within intragenic regions. The top transcription factors with binding motifs found in intergenic sites were CTCF, SIN3A, and RAD21. All three transcription factors are crucial in regulating chromatin structure to repress gene transcription and inhibition of these factors are closely associated with human disease (Davis et al. 2018; Witteveen et al. 2016; Zuin et al. 2014).

**Figure 1.**
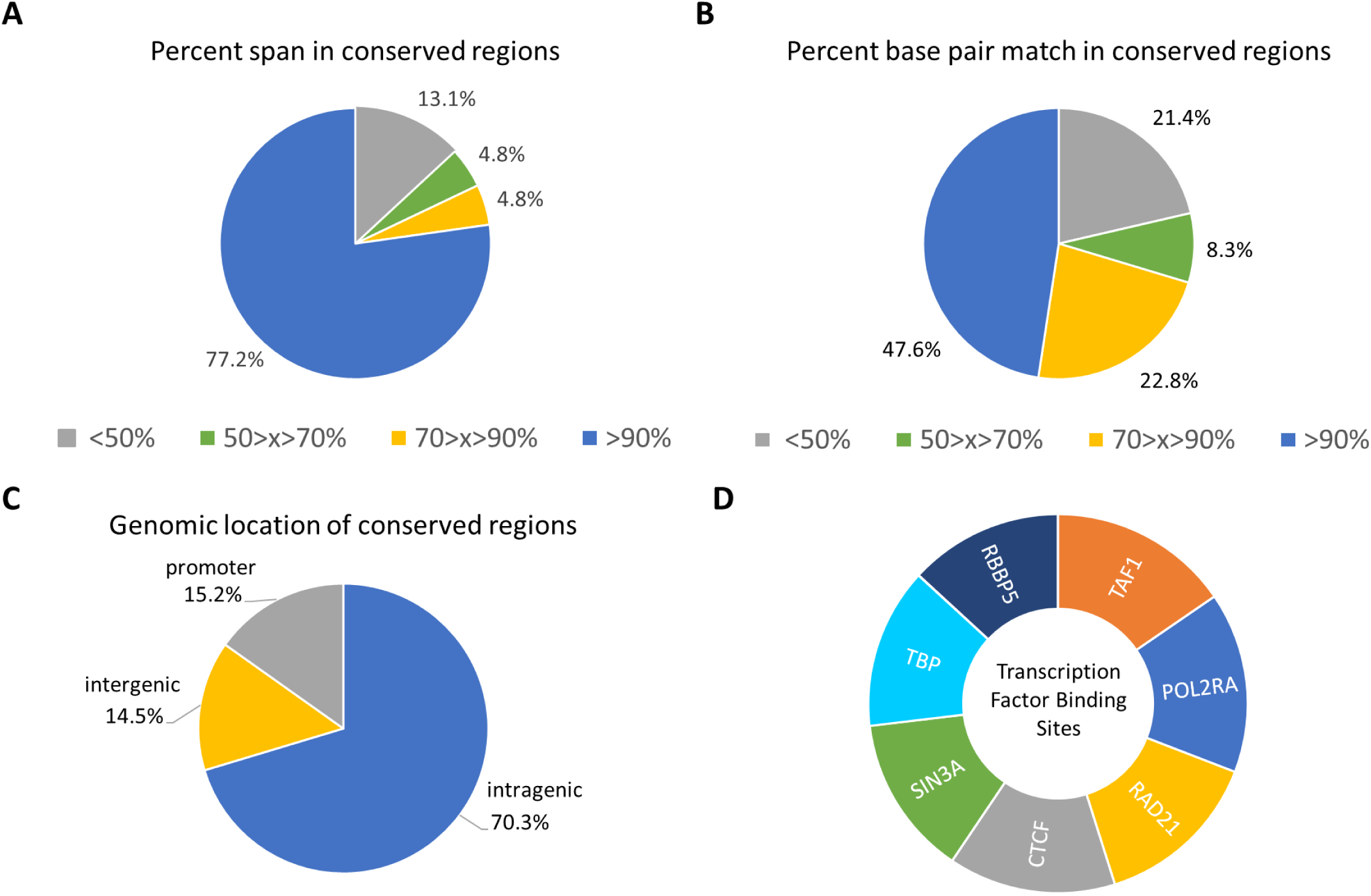
Characterization of DNMT1 dependent regions from mouse conserved in human genome. A) The percent span of the DNMT1 dependent regions in mouse covered by the identified human conserved regions. Pie chart represents the percentage of all identified conserved regions in the human genome that fall into each category. B) The percent base pair match of the DNMT1 dependent regions in mouse with the identified human conserved regions. Pie chart represents the percentage of all conserved regions in the human genome that fall into each category. C) The percentage of all conserved regions in the human genome that are located within the promoter, the gene body, or in non-coding intergenic regions. D) The top transcription factor binding sites found within all human conserved regions.

### Human DNMT1 dependent genes are enriched in cellular processes associated with cell-cell signaling

Out of the 145 regions identified in the human genome, we selected 97 of the most highly conserved regions compared to the mouse genome. The nearest human genes (112 genes) were used for functional enrichment analyses (Table 1). We observed that DNMT1 dependent genes were highly associated with cell to cell interactions and signaling (Figure 2A). Genes regulating cell-cell adhesion belonged primarily to the cadherin protein family. This agrees with previous reports showing monoallelic expression of protocadherins in Purkinje neurons (Esumi 2005). Gene Ontology of cellular components describe the subcellular compartments where enriched cellular processes and molecular functions occur. The plasma membrane and the pre-synapse were significantly enriched in our dataset in accordance with the enrichment of cell-cell interactions and calcium ion binding (FDR<0.05, Supplemental Table 3).

**Figure 2.**
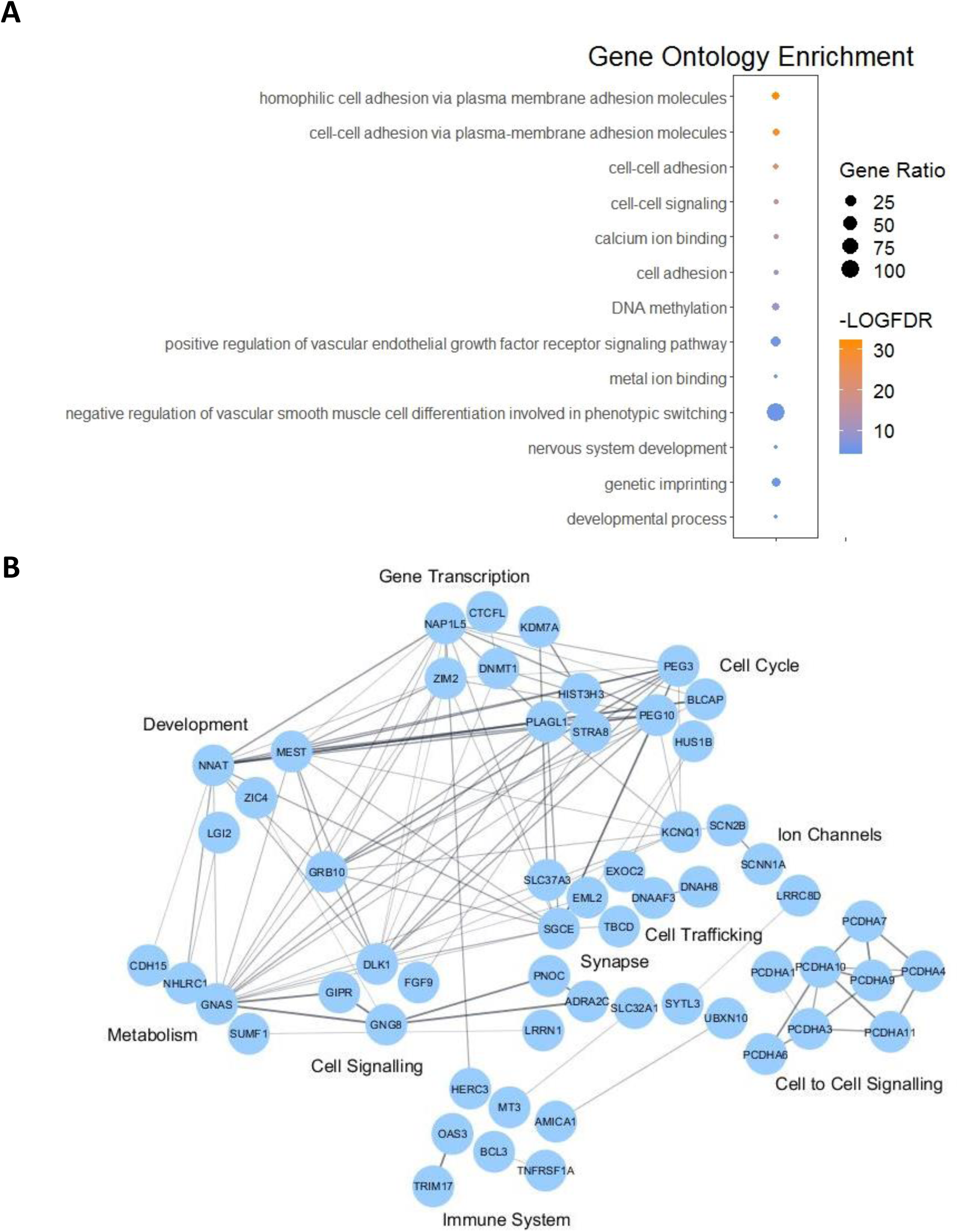
Functional enrichment analysis of DNMT1 dependent genes conserved in the human genome. A) Gene Ontology enrichment for cellular processes and molecular functions from the 112 selected human DNMT1 dependent genes (FDR<0.05). B) Network interactions from STRING database with an interaction enrichment p-value < 1x 10^−16^. Genes are clustered based on functional gene annotations from reactome pathways.

The interaction of the proteins encoded by DNMT1 dependent genes was analyzed using the STRING database (Figure 2B). The STRING database is a commonly used platform that summarizes the functional associations of a group of proteins. Out of the 112 selected protein encoding genes in humans, 98 nodes with 109 edges were detected with a medium confidence interaction score (>0.4). The interaction p-value (PPI) was less than 1×10^−16^, indicating that the number of associations was significant. We manually clustered genes with interactions using functional gene annotations. The largest cluster consisted of genes involved in cell-cell signaling including cadherins and cell surface adhesion molecules, cell trafficking chaperones, cytoskeleton proteins, and voltage-gated ion channels. Other significant pathways included developmental pathways of the nervous system and the vascular system.

### Human DNMT1 dependent genes are enriched in tissues of the brain

Given the evidence that DNMT1 dependent genes may play an important role in cell-cell adherence and communication as well as in nervous system development, we hypothesized that DNMT1 dependent genesm may be highly expressed in the brain. We analyzed the enrichment of tissue expression using the ARCHS4 human tissue database in EnrichR. The ARCHS4 database reports publicly available RNA-sequencing across all tissues and cell types in approximately 85,000 human samples (Lachmann et al. 2018). Significant expression of the DNMT1 dependent genes in the adult and developing brain was observed (p <0.01, Figure 3A). We also observed enrichment in the regions of the brain associated with motor function control. These include structures such as the cerebellum, spinal cord, and the striatum which are directly involved with motor coordination as well as the superior frontal gyrus which contains the supplementary motor area activated in complex movements (Li et al. 2013, Sang et al. 2015).

**Figure 3.**
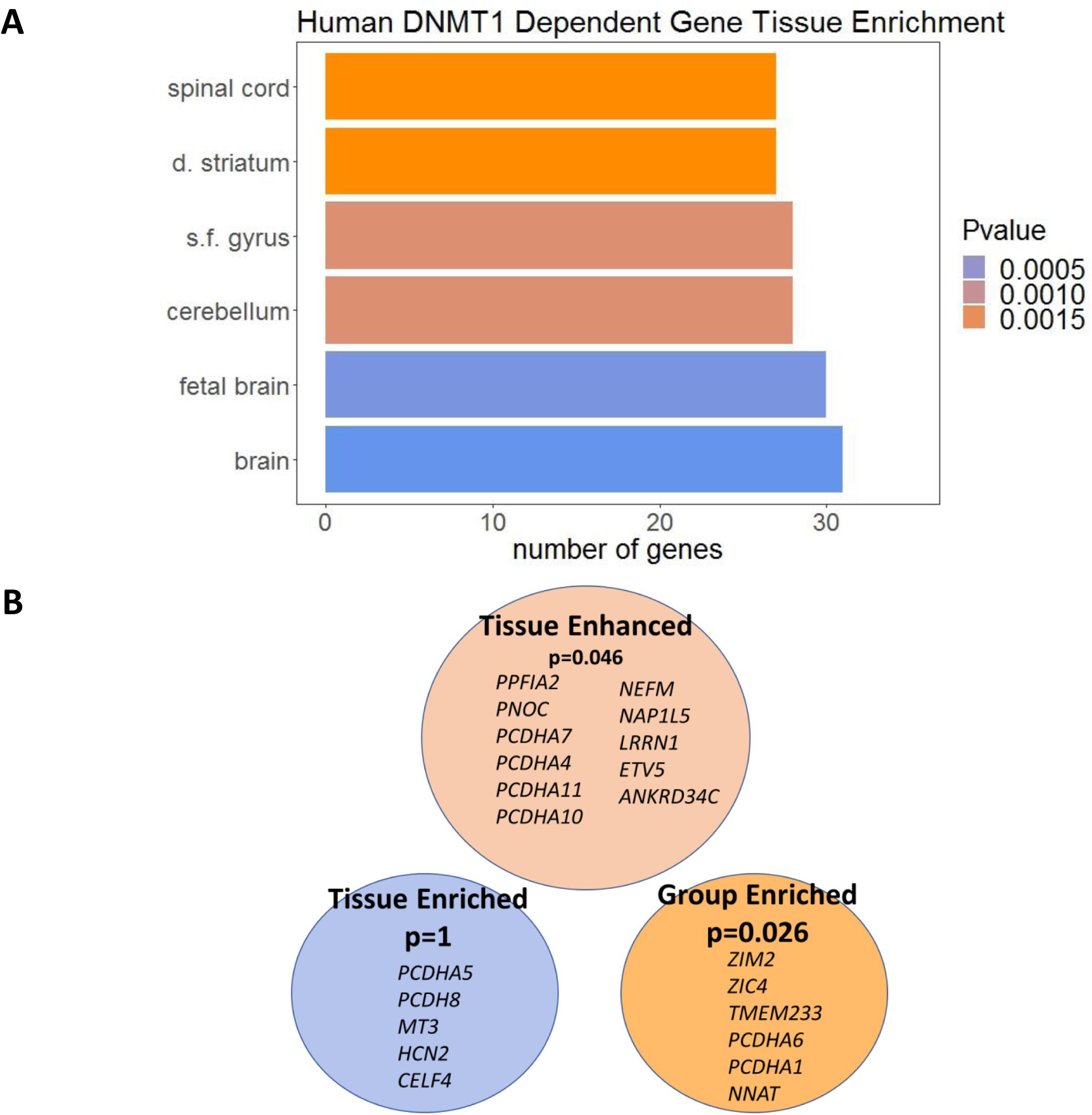
Tissue enrichment analysis of DNMT1 dependent genes conserved in the human genome. A) The top 6 tissues represented from the 112 selected human DNMT1 dependent genes (adjusted p-value<0.01) scored from Enrich R using the ArchS4 human tissue database (s.f. gyrus = superior frontal gyrus; d. striatum= dorsal striatum). B) DNMT1 human genes that are tissue-enriched, tissue-enhanced, or group-enriched genes within the cerebral cortex from Tissue Enrich (adjusted p-value shown).

To further determine if these genes were specific to neuronal tissues or if they have functionality across several tissues, we analyzed DNMT1 dependent genes for tissue specific enrichment using the TissueEnrich R package. Genes with increased expression in one tissue compared to all other tissues were defined as tissue enriched while genes with increased expression in one tissue compared to the average of all other tissues were defined as tissue enhanced. Group enriched genes were defined as genes that have increased expression in a group of tissues compared to all other tissues. Our analysis demonstrates that DNMT1 dependent genes conserved in the human genome have a significant abundance of tissue enhanced and group enriched genes within the brain, but not tissue enriched genes (Figure 3B). From these data, we conclude that DNMT1 dependent genes are likely important in cellular processes in the fetal and adult brain.

### Five DNMT1 dependent genes are represented in genes for potential PD blood biomarkers in patients

To further explore the significance of identified human DNMT1 dependent genes, we evaluated the recent literature on potential blood biomarkers in Parkinson’s disease (PD) patients. Encouragingly, we observed five differentially methylated genes in these studies within our conserved human regions (Henderson-Smith et al 2018; Wang et al. 2019). They are *COL9A2, SCNN1A, AMICA1, SLC16A3*, and *DLK1*. One of these genes, *COL9A2*, was also found to be differentially expressed in our rotenone treated cells (Henderson-Smith et al. 2018) (described below). Given that candidate PD biomarkers and our DNMT1 dependent genes were selected by differing criteria, we consider five overlapping genomic regions to be promising toward our hypothesis.

### Human DNMT1 dependent genes are differentially expressed in response to rotenone in human cells

We have shown that DNMT1 dependent regions have conserved sequences in the human genome and are enriched at genes involved in cell to cell interactions. These genes have enhanced expression in the brain and may contribute to neurological dysfunction and disease in response to environmental stress. To test the hypothesized contribution, we focus on rotenone exposure. Rotenone is a mitochondrial complex I inhibitor that is known to disrupt neuronal cell function in Parkinson’s disease associated brain regions (Tanner 2011). These brain regions include the cerebellum, spinal cord, striatum, and basal ganglia. We treated human cell line HEK293 with rotenone (200nm) for 24h. This dose was chosen based on previous reports in HEK293 and other neuronal cell models (Harris et al. 2018, Orth et al. 2003, Teixeria et al. 2018). Rotenone treatment had a substantial effect on the expression levels of over 2,000 genes (≥1.5 fold, FDR≤0.05). We examined these differentially expressed genes with our identified human DNMT1 dependent genes and discovered 14 of them had been changed upon rotenone treatment (Table 2). We validated the expression of 7 these genes with qRT-PCR (R^2^= 0.69, Supplemental Figure 2). We investigated whether these genes may contribute to rotenone-induced Parkinson’s disease by observing their expression in Parkinson’s disease tissues (Figure 4A). All 14 genes were expressed in Parkinson’s disease regions (>1 TPM) and 8 of the genes (*PPFIA4, NEFM, HCN2, ADRA2C, COL9A2, LRRC8D, EML2*, and *KDM7A/JHDM1D*) had pronounced expression in Parkinson’s disease regions (>35 TPM). Two genes, *NEFM* and *HCN2*, had significant expression (>100 TPM) in all selected regions and were identified by our tissue specific enrichment analysis as tissue enhanced and tissue enriched, respectively (Figure 3B). We therefore selected *NEFM* and *HCN2* for targeted methylation analysis based on their regional expression and significant up-regulation from both RNAseq and qRT-PCR analyses. The relevance and significance of *HCN2* and *NEFM* in human development and diseases are detailed later in the discussion.

**Table II.** Human DNMT1 dependent Genes Altered by rotenone.

| Gene | Log2FC | FDR | Region Type | Function/Process |
| --- | --- | --- | --- | --- |
| *MYH3* | -1.62 | 3.31E-09 | intragenic | myosin protein; cell movement and transport |
| *PPFIA4* | -1.37 | 2.59E-13 | intragenic | neurotransmitter release synaptic function |
| *COL9A2* | -0.79 | 1.53E-04 | intragenic | collagen; extracellular matrix organization |
| *DNAAF3* | -0.78 | 3.59E-02 | intragenic | dynein protein assembly; cell movement |
| *LRRC8D* | -0.78 | 8.42E-05 | intragenic | ion channel protein; neurotransmission |
| *PLEKHG4* | -0.65 | 2.74E-03 | intragenic | guanine exchange factor; cell signaling |
| *ADRA2C* | 0.59 | 2.93E-02 | intergenic | neurotransmitter release synaptic function |
| *EML2* | 0.62 | 5.99E-05 | promoter | microtubule protein; synaptic function |
| *HCN2* | 0.66 | 1.10E-03 | intragenic | voltage gated ion channel; action potential |
| *KDM7A* | 0.67 | 1.16E-04 | intergenic | histone demethylase; neurodevelopment |
| *PHLDB3* | 0.76 | 3.75E-04 | intragenic | enzyme binding; cell growth and proliferation |
| *GIPR* | 0.76 | 2.37E-02 | promoter | gastric inhibitory peptide; insulin release |
| *BCL3* | 0.78 | 2.00E-02 | intragenic | proto-oncogene; cell growth and proliferation |
| *NEFM* | 0.80 | 4.33E-07 | intragenic | neurofilament; synaptic function |

**Figure 4.**
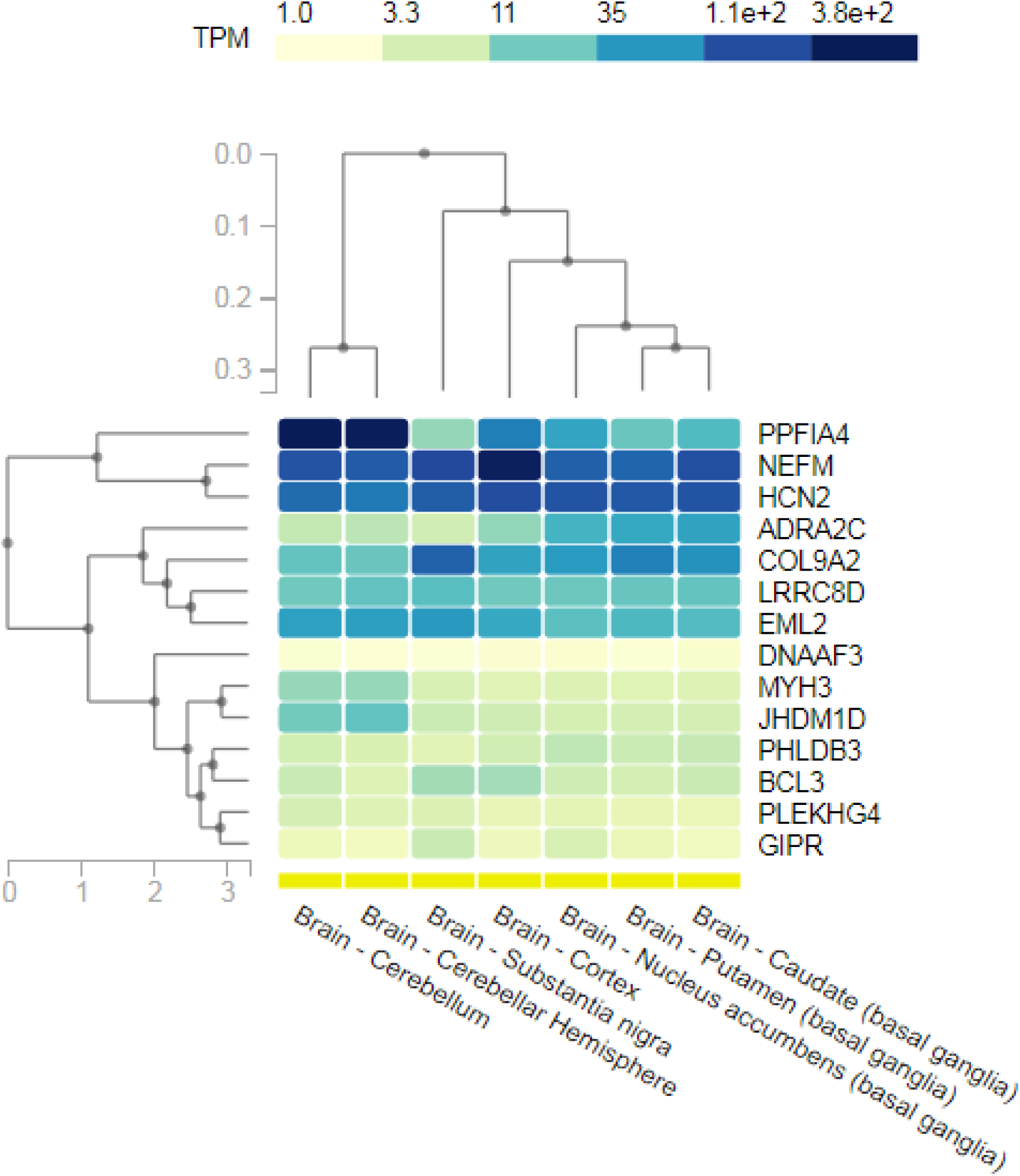
Regional expression of DNMT1 dependent genes altered by rotenone. Human DNMT1 dependent genes that were differentially expressed (≥1.5 fold change, FDR≤0.05) in response to rotenone were used for regional expression analysis in the Genotype-Tissue Expression (GTex) database. Brain regions selected are associated with Parkinson’s disease pathogenesis and important for motor function control. The heat map was generated using GTex and organization is clustered by gene function and tissue function. The color of each square indicates the level of expression of the gene in the selected region in transcripts per kilobase million (TPM).

### DNMT1 dependent regions at *HCN2* and *NEFM* are differentially methylated in response to rotenone in human cells

Previously, germline ASMs are especially vulnerable to environmental exposure, thereby altering imprinted gene expression (Susiarjo et al. 2013). Herein, we determined the potential methylation changes of the defined DNMT1 dependent region at these two gens, *NEFM* and *HCN2*, with significant up-regulation in response to rotenone. We completed base-resolution bisulfite sequencing of these regions amplified with bisulfite PCR. After filtering of low-quality reads, approximately 42% of reads were mapped uniquely to the amplified regions. We observed high correlation between biological replicates and similar average coverage between control and treated samples (Supplemental Figures 2-3). The average CpG coverage for both genes in all samples was >15,000x. Of the 23 predicted CpG sites within the amplified DNMT1 dependent region on exon 8 of *HCN2*, 21 CpGs had adequate coverage (>1000x) in all samples and 14 CpGs were differentially methylated. These differentially methylated cytosines were largely hypomethylated (Supplemental Figure 4). The mean percent methylation of all CpGs within the amplified DNMT1 dependent region was also significantly hypomethylated (Δme of -1.84%, FDR<0.05). We saw a similar trend in the DNMT1 dependent region at exon 1 of *NEFM*. Of the 39 predicted CpG sites within the amplified DNMT1 dependent region, 35 CpGs had adequate coverage in all samples and 13 of these CpGs were differentially methylated. A slight majority (54%) of these differentially methylated cytosines were hypomethylated (Supplemental Figure 4). The overall change in methylation ratio for the entire region was significant but very low (<0.1% absolute difference, FDR<0.01). As a result, we decided to focus on the first 200 of the 500 bp amplified region, which overlap both the CpG island at exon 1 as well as a CTCF transcription factor binding site reported by ENCODE (ENCODE 2012). In this region, there was a slightly higher change in methylation (Δme of -0.12%, FDR<0.05). Additionally, we used the CTCF binding site prediction tool to determine the exact CpG sites within the CTCF binding motif (Ziebarth and Bhattacharya 2013). One of the top hits predicted CpG binding on the negative strand at CpG sites 89-96 within the defined *NEFM* region (Figure 6; Supplemental Table 5). Three of these four CpG sites were differentially methylated with half of them having >2% reduced methylation.

**Figure 5.**
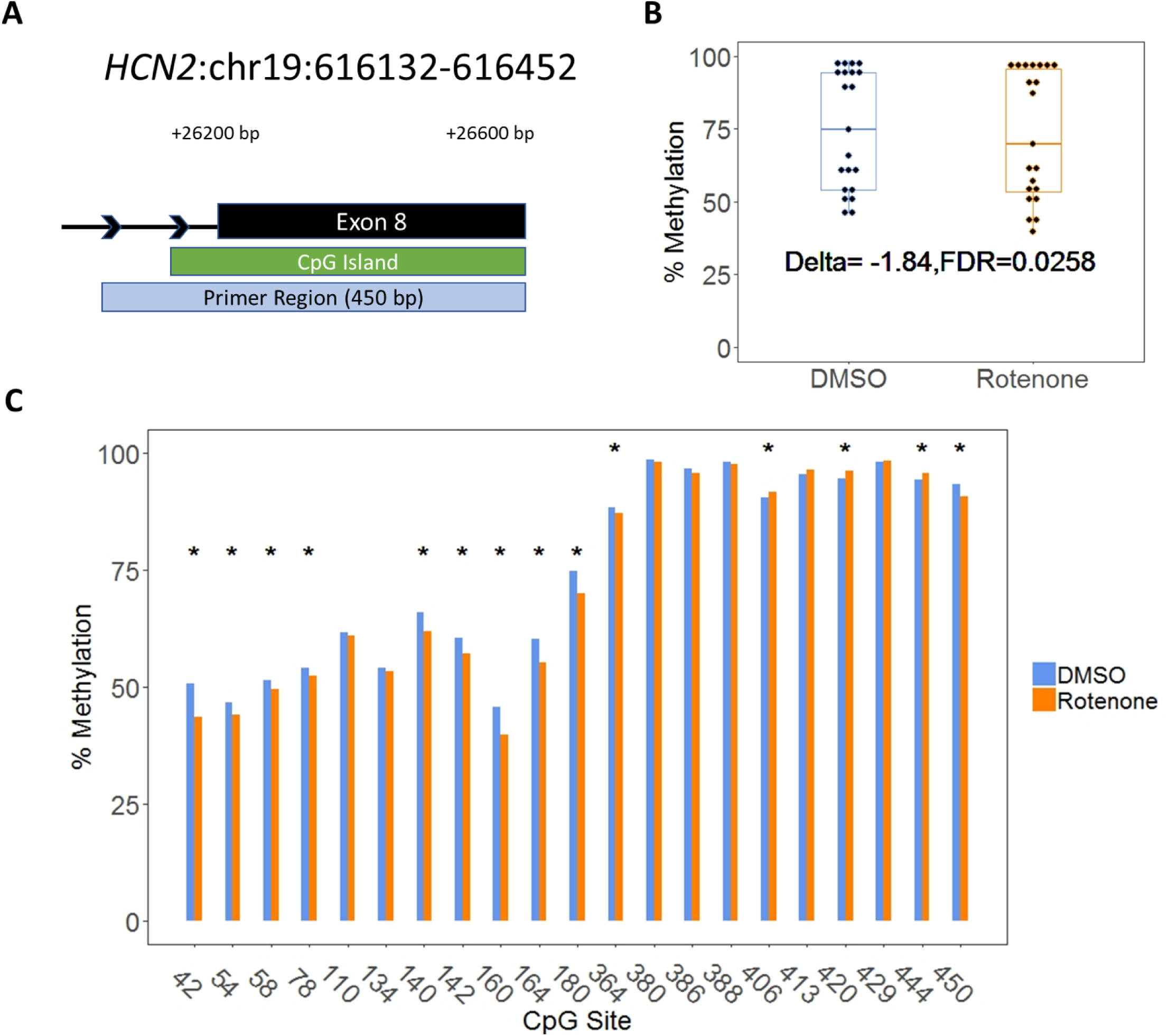
Altered CpG methylation at *HCN2* human DNMT1 dependent locus. A) Genomic location of identified *HCN2* DNMT1 dependent region. The DNA element and distance from the transcription start site is annotated in black. The primer region box indicates the amplified region for Bisulfite-sequencing. B) The percent methylation of all CpG sites within the amplified region. Delta indicates the change in the mean CpG methylation percentage and the associated false discovery rate. C) The percent methylation of individual CpG sites within the amplified region. Significant differentially methylated cytosines are indicated by * (Δ>1%; q-value<0.01).

**Figure 6.**
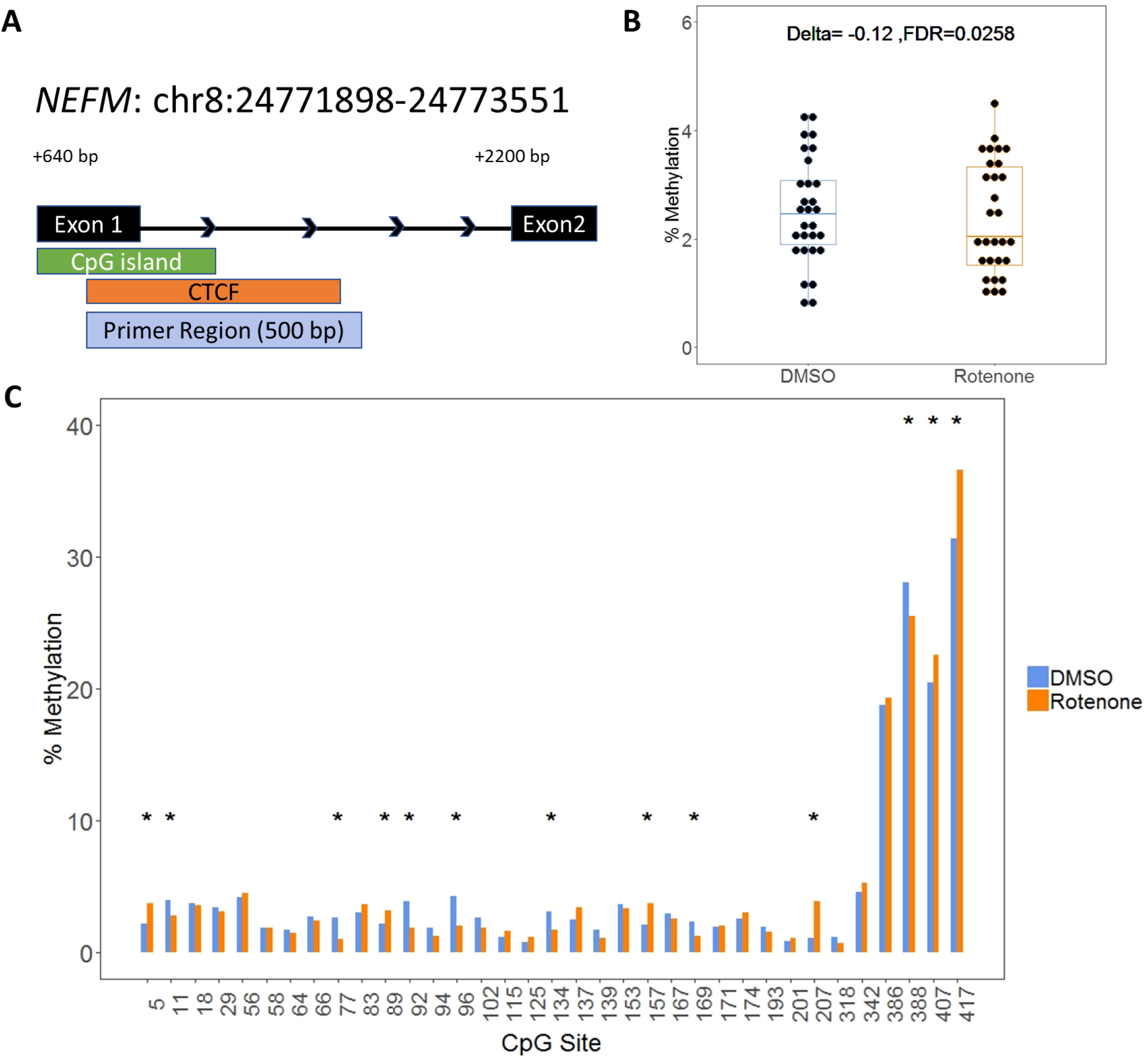
Altered CpG methylation at *NEFM* human DNMT1 dependent locus. A) Genomic location of identified *NEFM* DNMT1 dependent region. The DNA element and distance from the transcription start site is annotated in black. The transcription factor binding site for CTCF was annotated from ENCODE v2 and ENCODE Uniform TFBS tracks in Genome Browser. The primer region box indicates the amplified region for Bisulfite-sequencing. B) The percent methylation of all CpG sites within the first 200 base pairs of the amplified region. Delta indicates the change in the mean CpG methylation percentage and the associated false discovery rate. C) The percent methylation of individual CpG sites within the amplified region. Significant differentially methylated cytosines are indicated by * (Δ>1%; q-value<0.01).

These data suggest that the methylation status of DNMT1 dependent regions in the human genome are vulnerable to the neurotoxicant rotenone. We found that the coding regions and transcription factor binding motifs may be among the DNA elements that are particularly susceptible to exposure. The changes in methylation we observed were similar in scale to observed differential methylation at gene-encoding regions in the blood and brain of Parkinson’s disease patients (Masliah et al. 2013; Navarro-Sanchez et al. 2018; Henderson-Smith et al. 2018; Wang et al. 2019). Both *HCN2* and *NEFM* are regionally expressed in Parkinson’s disease tissues and their function has a critical role in neuronal plasticity and survival detailed in the discussion.

## Discussion

Our previous work identified DNMT1 dependent putative non-germline ASMs in the mouse genome. In this study, we analyzed these regions and found that 70% were in highly conserved regions with the human genome. In the human genome, our candidate loci were often located at gene-coding regions and half of them overlapped transcription factor binding sites (Figure 1). Methylated cytosines alter gene expression by influencing the binding of transcription factors to DNA. We listed the top ten transcription factor binding sites within our identified candidate regions in human embryonic stem cells using ENCODE experimental data (Figure 1). Unsurprisingly, three of these transcription factors were TAF1, TBP, and POL2RA which all have an essential role in initializing transcription. We were interested to see SIN3A and RBBP5 which both interact with histone modifying enzymes to regulate chromatin accessibility and are critical during neurodevelopment (Gabriele et al. 2018). Furthermore, SIN3A is recruited to the methyl-CpG binding protein MeCP2 to silence transcription. Mutations in MeCP2 cause an X-linked neurodevelopmental disorder known as Rett Syndrome and similarly impairment of SIN3A expression also causes developmental cognitive deficits (Witteveen et al. 2016). MeCP2 and SIN3A have been linked to the establishment and maintenance of imprinting control regions but their effect on the expression of neighboring imprinted genes remains to be determined (Ma et al. 2015).

CTCF is another transcription factor of interest with 17% of the DNMT1 dependent human regions overlapping the CCCTC-binding motif. CTCF is also critical in neurodevelopment and chromatin organization (Franco et al. 2014, Davis et al. 2018). CTCF mediates the formation of chromatin loops and thus can promote widespread changes in gene expression (Hou et al. 2008, Phillips and Corces 2009). When bound to sequences known as insulator sequences, CTCF represses transcription by blockading promoter-enhancer interactions (Bell et al. 2000, Hark et al. 2000). CTCF and the stabilizing protein cohesion bind at numerous imprinted control regions (Rubio et al. 2008, Prickett et al. 2013). CTCF has been reported to preferentially bind unmethylated chromatin but binding affinity depends not only on the methylation status of the motif itself but the surrounding CpG sites as well (Li et al. 2017, Wang et al. 2012).

Given the importance of imprinted gene clusters in development, we analyzed functional enrichment of our non-germline ASMs in the human genome. The most significant biological process associated with our gene set was cell to cell adhesion (Figure 2). We observed a significant group of cadherins encoded adjacent to DNMT1 dependent regions on chromosome 5. Cadherin proteins are expressed on the membrane of embryonic stem cells and are critical for their self-renewal by forming tight intracellular niches (Pieters and Frans van Roy 2014). The expression of cadherin subtypes on embryonic stem cells is variable and the patterning of cadherin expression also controls their differentiation. Protocadherins are involved in neuronal connectivity and this function extends from neural progenitors during development into postmitotic neurons in the adult brain (Sams et al. 2016). Intriguingly, protocadherin is regulated by CTCF and deletion of CTCF in mice caused deficits in hippocampal learning and memory via dysregulation of protocadherin expression (Sams et al. 2016). The most significant molecular function was calcium ion binding and the pre-synaptic axon terminal was one of two most significant cellular components represented. This agreed with our network analyses where multiple genes were involved in cell trafficking and synaptic activity (Figure 2). We investigated whether developmental genes were specific to an individual tissue or group of tissues. These genes from DNMT1 dependent regions have significant enrichment of genes expressed in the cerebral cortex from two separate databases, EnrichR ArchS4 and Tissue Enrich Human Protein Atlas (Figure 3). Our data suggests that DNMT1 dependent non-germline ASMs from the human have enhanced expression in the brain, which could be important for neurological development and cellular communication function.

In our previous work, we characterized non-germline ASMs in the mouse genome at two genes, *HCN2* and *Park7*, with potential in Parkinson’s disease (Bonifati et al. 2003, Kim et al. 2005, Martos et al. 2017). The proper maintenance of the epigenome throughout aging is believed to have a major impact on the risk of neurodegeneration later in life (Gapp et al. 2014, Labbe et al. 2016). The influence of germline-ASMs on neurodegeneration has recently been of interest in the literature given their involvement in neurodevelopment but the effect of non-imprinted ASMs have not been well characterized (Gapp et al. 2014). To experimentally examine the association of identified ASMs in the human genome with Parkinson’s disease, we used human embryonic kidney cells with a neuronal lineage phenotype and treated them with rotenone for 24h (Stepanenko and Dmitrenko 2015). We observed several of our candidate genes were affected in response to rotenone treatment and half of these genes have regional expression in Parkinson’s disease associated regions (Figure 4). Among these genes, *HCN2* and *NEFM* were determined from our tissue enrichment analysis to have a higher expression level in the brain than any other tissue. Additionally, experimental analysis of *HCN2* in the mouse genome suggests a random, switchable allele-specific methylation pattern that was independent of the parent-of-origin (Martos et al. 2017). We selected these two genes for methylation analysis to determine if conserved non-germline ASMs in the human genome were sensitive to environmental factors associated with Parkinson’s disease.

The *HCN2* gene encodes an isoform of the hyperpolarization-activated cyclic nucleotide-gated channel located on the membrane of neurons in the central and peripheral nervous system. HCN channels regulate neuronal plasticity and have the advantage of using both voltage dependent mechanisms as well as cAMP intracellular signaling mechanisms (DiFrancesco et al. 1999). In the midbrain, these channels control the spontaneous activity of dopaminergic neurons and their dysfunction has been linked to the depletion of dopamine in Parkinson’s disease (Chen et al. 2011, DiFrancesco and DiFrancesco 2015, Good et al. 2011). In the human genome (hg19), the conserved DNMT1 dependent locus identified was 321 bp at a CpG island on exon 8 of the gene. We observed significant upregulation of mRNA expression levels (1.6 fold change, FDR<0.01) that correlated with DNA hypomethylation (−1.8%, FDR<0.05) of a 450 bp site surrounding the region of interest (Table 2; Figure 5; Supplemental Table 4). Dysregulated *HCN2* expression could affect HCN2 channel activity leading to disrupted regulation of dopaminergic excitability.

The *NEFM* gene encodes a subunit of neuron-specific intermediate filaments known as neurofilaments. Neurofilaments are primary components of myelinated axons and are essential for synaptic function (Yuan et al. 2017). Neurofilament subunit expression is tightly regulated to maintain proper stoichiometry. As such, aberrant expression of *NEFM* likely disrupts axonal growth and transport. Interestingly, neurofilament subunits including the NEFM protein are considered promising neurodegeneration biomarkers due to their cell specificity and sensitivity to neuronal damage (Khalil et al. 2018). In Parkinson’s disease patients, neurofilament proteins have been detected at higher levels in the cerebral spinal fluid, and more recently, in the blood (Abdo et al. 2007, Rosengren et al. 1996, Rojas et al. 2016). In our data, the conserved DNMT1 dependent locus covered a 150 bp region in exon 1 as well as a 1,650 bp region spanning exon 1 to intron 2. We observed significant upregulation of mRNA (1.7 fold change, FDR<0.01) and hypomethylation of a 200 bp section of the identified region at exon 1 (−0.12%, FDR<0.05). While the total change in percent methylation was relatively small, the selected region contained a CTCF binding site. Several CpG sites located within this CTCF binding motif had higher changes in methylation (>1%, adjusted q-value<0.01) (Figure 6; Supplemental Table5). As mentioned previously, CTCF is hypersensitive to changes in DNA methylation and approximately 41% of CTCF binding variability has been attributed to DNA methylation (Wang et al. 2012). The lack of repressive signaling from CTCF could contribute to the observed increases in NEFM reported in Parkinson’s disease patients and the observed overexpression of *NEFM* associated with cytoplasmic inclusions in motor-impaired mice (Sosa et al. 2003, Liu et al. 2011, Wong et al. 1995).

DNMT1 expression in neural stem cells is essential for adult neurogenesis and the survival of adult neurons in the brain (Noguchi et al. 2015). We’ve shown that non-germline ASMs are dependent on DNMT1 in mice. The goal of this study was to identify conserved DNMT1 dependent regions and putative non-germline ASMs in the human genome and test the vulnerability of these regions to a neurotoxicant associated with Parkinson’s disease. Our work identified candidate, non-germline ASMs with DNMT1 dependence as enriched in the human brain. We discovered 14 genes have altered expression (>1.5 fold change) at predicted ASMs in response to rotenone.

We evaluated the recent literature on candidate blood biomarkers in Parkinson’s disease patients and observed five differentially methylated genes (*COL9A2, SCNN1A, AMICA1, SLC16A3*, and *DLK1*) in these studies within our conserved human regions (Henderson-Smith et al 2018; Wang et al. 2019). Of these genes, *COL9A2* was differentially expressed in our rotenone treated cells and was determined to have high regional expression in the substantia nigra (Table II, Figure 4) (Henderson-Smith et al. 2018). This observation is strengthened with another study that has found that differentially methylated genes in the blood have high concordance with differentially methylated genes in the brain (Masliah et al. 2013). These data partially support our hypothesis that environmentally induced changes in DNMT1 dependent ASMs in the human brain can alter the risk of neurodegeneration. To further examine our hypothesis, we quantified methylation of identified regions at 2 selected genes known to increase the risk for Parkinson’s disease and observed significant hypomethylation. In the future, a larger panel of these identified regions in the human genome will be tested in other cells lines at varying points in neuronal differentiation to determine the role of non-germline ASMs on neuronal development and its maintenance with age.

## Acknowledgements

This project is made possible with support from the JHU Catalyst Award (ZW), as well as student Frank Bang Award (to DMF). Experimental disposals and salaries were also partially supported by the U.S. National Institutes of Health (R01ES25761, U01ES026721 Opportunity Fund, and R21ES028351) to ZW and the National Institutes of Health T32 Pre-doctoral training grant (5T32ES7141).

## Data Availability

The data used to generate this report are available upon request from the corresponding author.

