## Supplementary for "The conserved DNMT1 dependent methylation regions in human cells are vulnerable to environmental rotenone"

**Supplementary Table 1. q-RTPCR primers for RNAseq validation in human genome**

| | |
|---|---|
| PPFIA4_For | CTCTGCGGATGTTGTCTCCC |
| PPFIA4_Rev | ATGCTGCCACTGGTTACACG |
| DNAAF3-For | GGGCTCAAGTCATTCACCCC |
| DNAAF3-Rev | GGTTGGGCACATGATAGGC |
| LRRC8D-For | ATGACATTCAGCCAACTTACCG |
| LRRC8D-Rev | TACTGGCAAACAGACCACCTG |
| ADRA2C-For | GCCTCAACGACGAGACCTG |
| ADRA2C-Rev | CCCAGCCCGTTTTCGGTAG |
| EML2-For | CTCCGTAGCCGTGCTATACAG |
| EML2-Rev | CCAAGCATTTGATGTCATCGTTG |
| HCN2-For | AGAAGGGCATTGACTCCGAG |
| HCN2-Rev | TAGCGGATCAGGCGTGAGA |
| NEFM-For | GCTCGTCATTTGCGCGAATAC |
| NEFM-Rev | TTTCTGTACGCAGCGATTTCTAT |
| GAPDH-For | ACAACTTTGGTATCGTGGAAGG |
| GAPDH-Rev | GCCATCACGCCACAGTTTC |

**Supplementary Table 2. BS-PCR primers for human genome**

| | |
|---|---|
| HCN2-For | TGAGATTATTTTGGTTAATATGGTGAA |
| HCN2-Rev | CAAAAAAATAAAATTCTTCTTACCTAC |
| NEFM-For | GGTATTAAGGAGTTTTTGGAG |
| NEFM-Rev | CTAACCTAACCATTCCCATCTAAAC |

**Supplementary Table 3. Gene Ontology Results for Cellular Component**

| GO Term | Cellular Component | Gene Ratio | FDR |
|---|---|---|---|
| GO:0005886 | plasma membrane | 0.008 | 0.024 |
| GO:0098793 | presynapse | 0.023 | 0.028 |

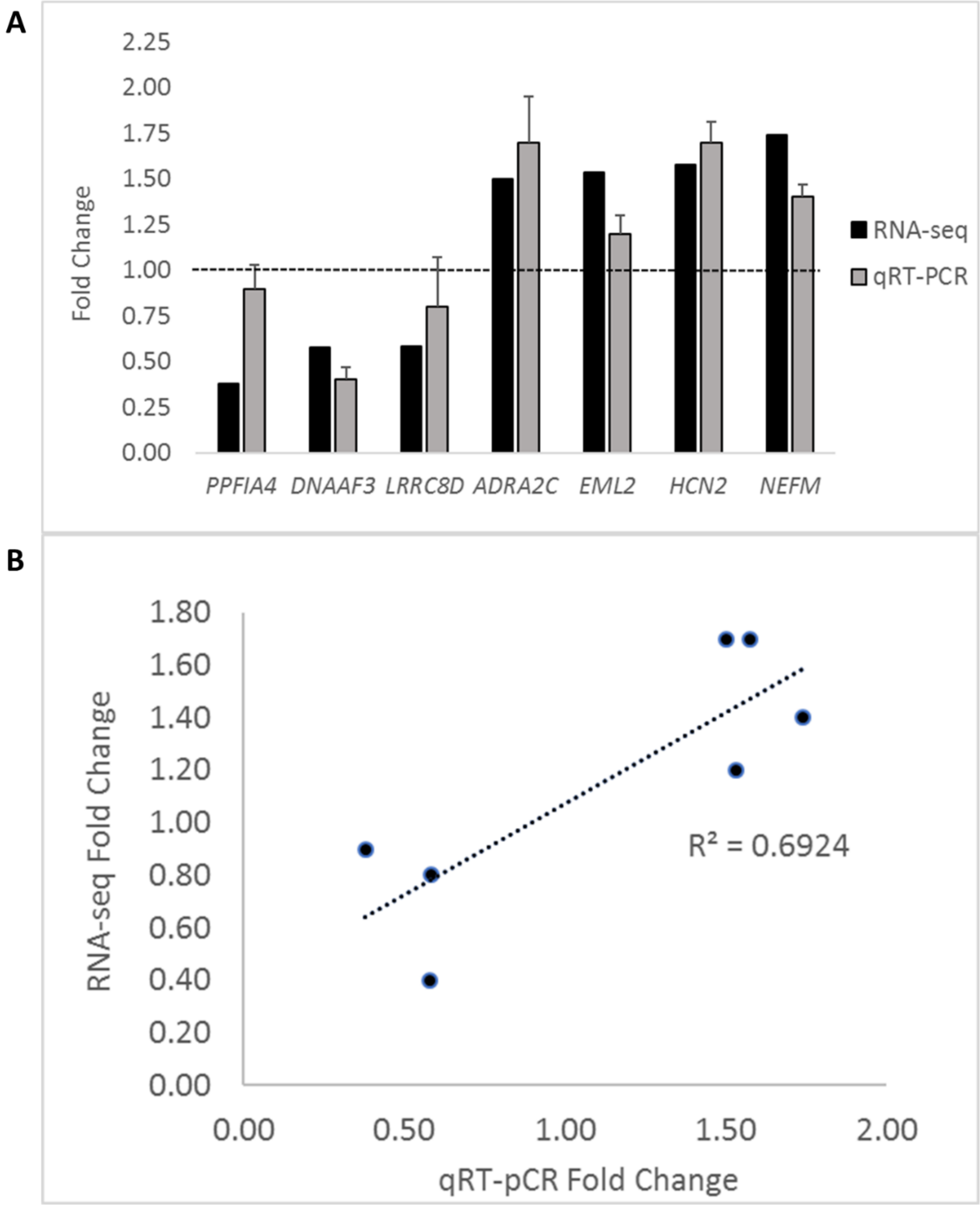


**Supplementary Figure 1. RNA sequencing validation with q-RTPCR.** A) Fold change comparison of RNA sequencing results versus q-RTPCR results. B) Linear calibration curve of RNA sequencing results with q-RTPCR results expressed as fold change in expression.

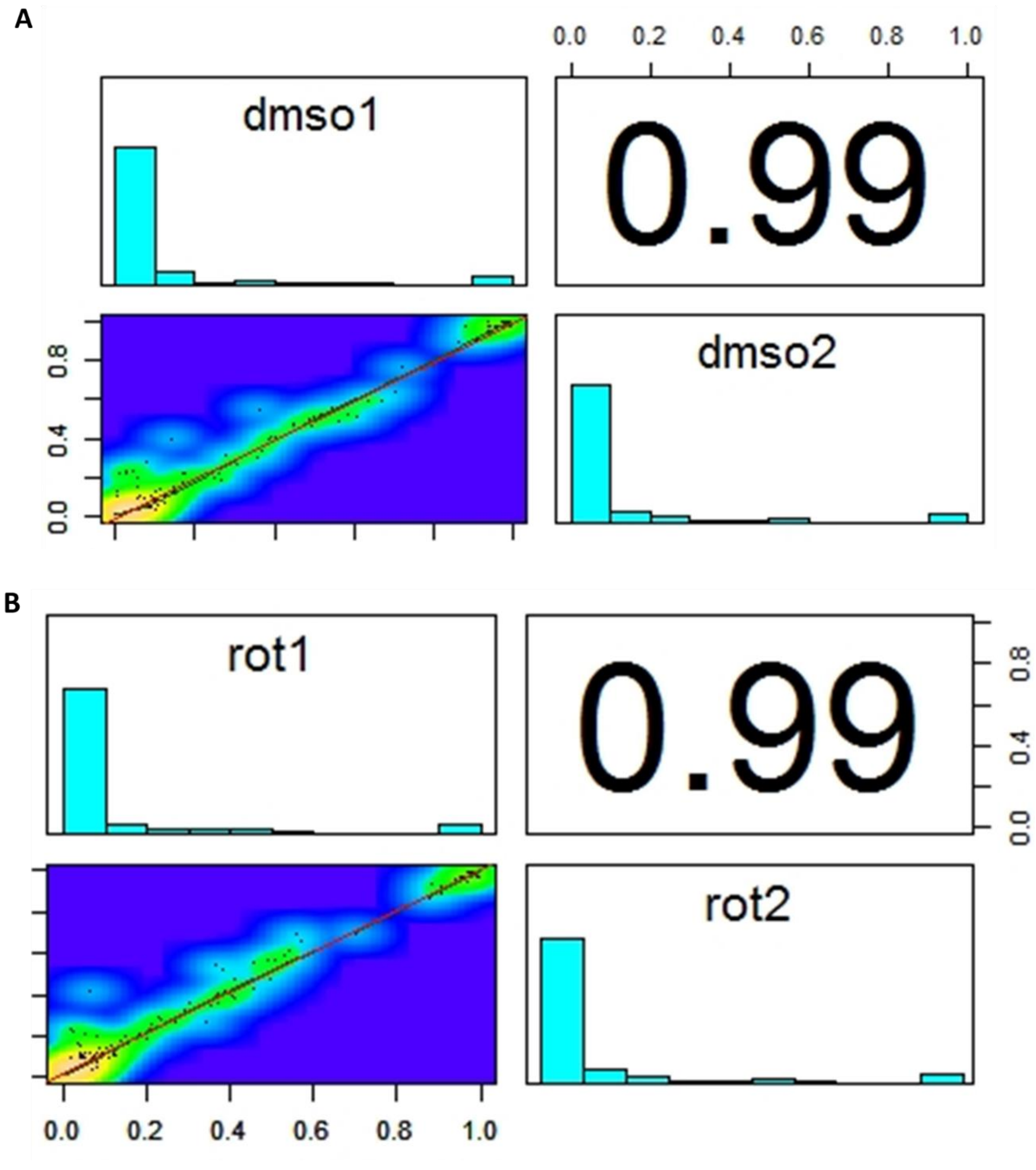


**Supplementary Figure 2. Pearson's correlation coefficient for bisulfite sequencing replicates of HEK293.** A) Correlation coefficients of Bisulfite sequencing data between biological replicates of control samples. B) Correlation coefficients of Bisulfite sequencing data between biological replicates of rotenone treated samples.

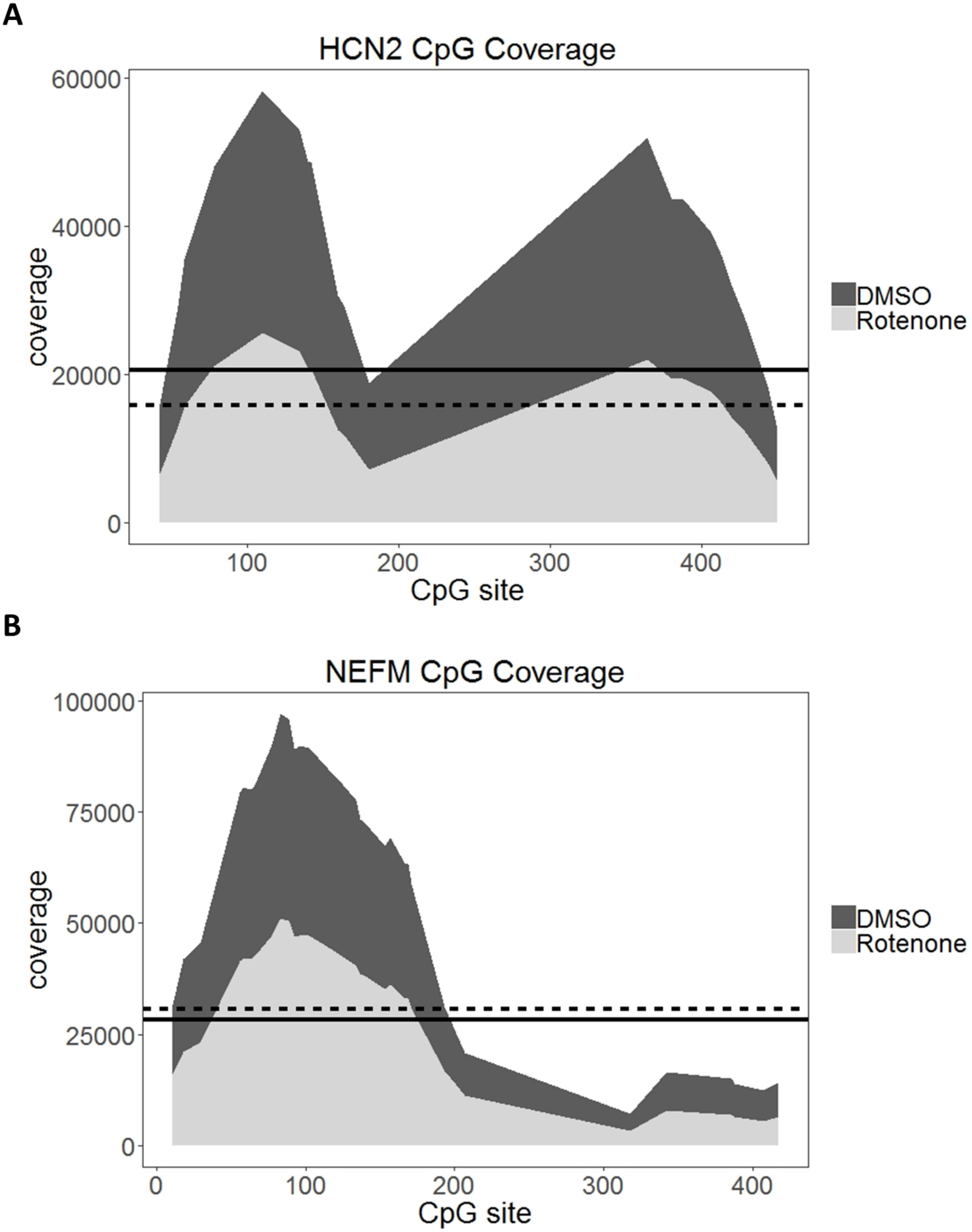
A
HCN2 CpG Coverage
60000
40000
20000
0
coverage
100
200
300
400
CpG site
DMSO
Rotenone
B
NEFM CpG Coverage
100000
75000
50000
25000
0
coverage
0
100
200
300
400
CpG site
DMSO
Rotenone

**Supplementary Figure 3. Bisulfite sequencing coverage of CpG sites within amplified DNMT1 dependent regions at *HCN2* and *NEFM*.** The average total coverage for all CpG sites within the amplified region is indicated by the straight line for DMSO and the dash line for rotenone.

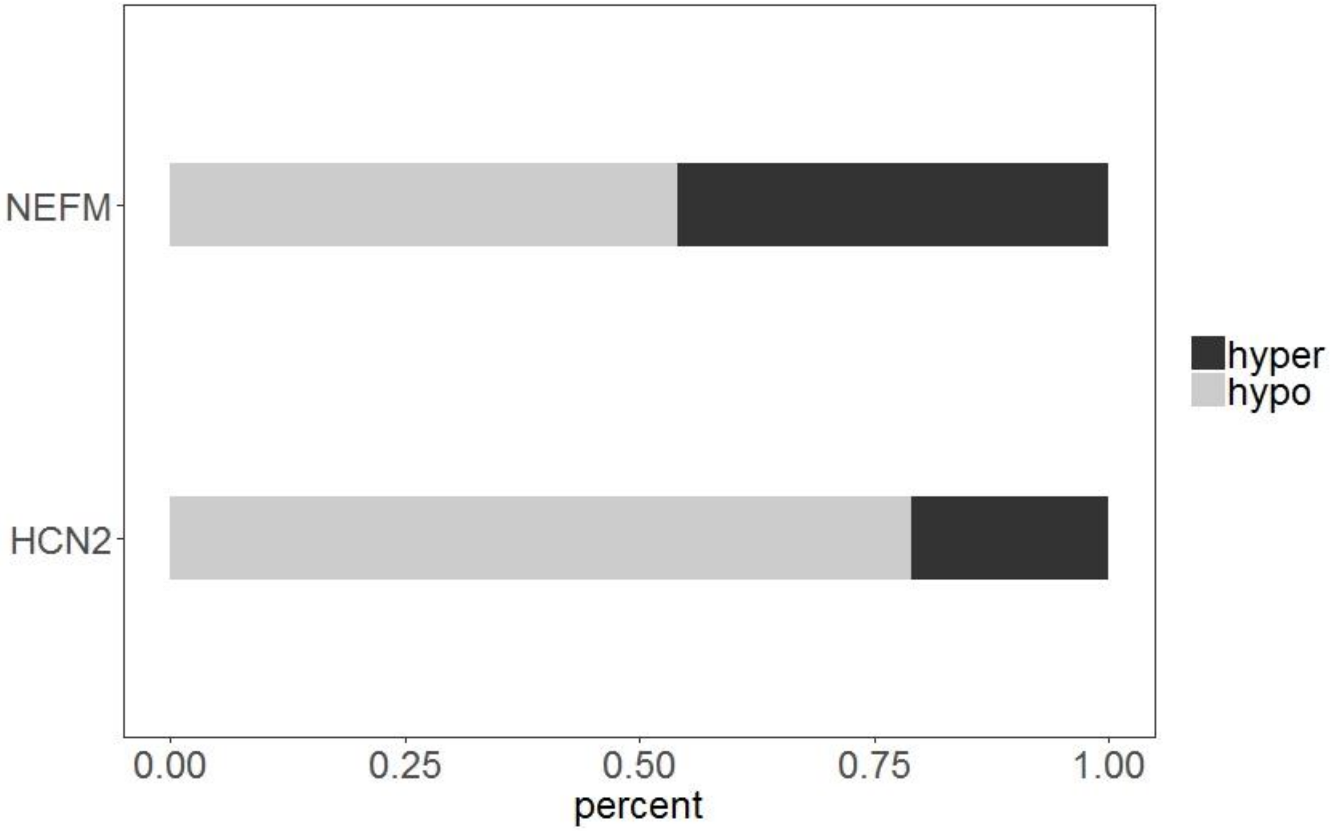


**Supplementary Figure 4. Percentage of hyper and hypo differentially methylated CpGs within DNMT1 dependent loci.**

**Supplementary Table 4. CPG Methylation HCN2**

| CpG site | q-value | Δme |
|---|---|---|
| 42 | 6.8E-18 | -7.0 |
| 54 | 5.4E-06 | -2.6 |
| 58 | 3.3E-04 | -1.8 |
| 78 | 1.7E-04 | -1.7 |
| 140 | 8.3E-19 | -3.9 |
| 142 | 3.6E-13 | -3.3 |
| 160 | 2.5E-25 | -6.0 |
| 164 | 7.6E-17 | -5.0 |
| 180 | 3.0E-13 | -4.9 |
| 364 | 5.0E-05 | -1.2 |
| 406 | 2.3E-04 | 1.0 |
| 420 | 1.2E-12 | 1.7 |
| 444 | 5.4E-06 | 1.5 |
| 450 | 2.1E-08 | -2.7 |

**Supplementary Table 5. CPG Methylation NEFM**

| CpG site | q-value | Δme |
|---|---|---|
| 5 | 6.0E-10 | 1.6 |
| 11 | 3.2E-09 | -1.2 |
| 77 | 7.8E-78 | -1.6 |
| 89 | 7.6E-24 | 1.1 |
| 92 | 8.6E-75 | -2.0 |
| 96 | 2.4E-83 | -2.3 |
| 134 | 3.4E-36 | -1.4 |
| 157 | 2.7E-34 | 1.6 |
| 169 | 2.9E-22 | -1.0 |
| 207 | 7.7E-37 | 2.7 |
| 388 | 4.0E-04 | -2.6 |
| 407 | 2.7E-03 | 2.1 |
| 417 | 9.5E-11 | 5.2 |
